# A Fundamental Relationship between TCR Diversity, Repertoire Size and Systemic Clonal Expansion: Insights from 30,000 TCR*β* Repertoires

**DOI:** 10.1101/2025.05.06.652462

**Authors:** H. Jabran Zahid, Damon May, Harlan Robins, Julia Greissl

## Abstract

TCR diversity is essential for immune defense, yet the mechanisms underlying its decline with age, its dependence on sex and its variation among individuals remain poorly understood. These patterns are often attributed to passive loss from factors such as thymic atrophy and cumulative immune exposures but such processes fail to explain the systematic variation observed across populations. Here we challenge this view by analyzing TCR*β* repertoires from *∼*30, 000 individuals showing that TCR diversity is almost entirely determined by repertoire size and the frequency of the 1,000 most abundant clones. These two intrinsic features of the repertoire explain 96% of the variance in TCR diversity, capturing its dependence on age and sex and defining a robust relationship that holds even under strong immune perturbations such as Cytomegalovirus infection. This relationship arises because the frequency of abundant clones captures a repertoire-wide pattern of coordinated clonal expansion—termed intrinsic clonality—which may be a fundamental, previously unrecognized property of the immune system. We propose that TCR diversity emerges as a system-level property mediated by repertoire size and intrinsic clonality, both of which are likely homeostatically regulated. These findings offer a new conceptual framework for understanding TCR diversity within immune homeostasis which may guide therapies aimed at restoring immune function.

## Introduction

The theory of clonal selection is a cornerstone of modern immunology, providing the foundation for understanding how TCR diversity is broadly shaped and maintained by the immune system (*1*). T cells play an essential role in immune defense by targeting antigens from infections and cancer, with their specificity determined by T cell receptors (TCRs) (*2, 3*). The ability to respond to a broad range of pathogens is enabled by the diversity of the TCR repertoire (*4*– *13*). A large pool of naive T cells with diverse, randomly rearranged TCRs is primarily generated during childhood and adolescence via V(D)J recombination and is maintained throughout adulthood predominantly through homeostatic proliferation (*14, 15*). Mechanisms of immune tolerance eliminate or regulate self-reactive T cells, thereby limiting responses to non-self antigens (*16*). The T cell repertoire is further shaped by selection processes, including clonal expansion upon antigen encounter and the subsequent preferential retention of activated T cells in the memory compartment. A key feature of immune homeostasis is the long-term balance between naive and memory T cell compartments, which enables rapid responses to previously encountered antigens while preserving the capacity to recognize new ones (*17, 18*).

Despite its essential role, the mechanisms governing TCR diversity remain poorly understood. Diversity declines by about a factor of two between the ages of 20 and 80 years and is systematically lower in males than females (*19, 20*). Moreover, variation between individuals exceeds that explained by age and sex alone (*20*). Changes in TCR diversity are commonly attributed to passive, cumulative processes such as thymic atrophy, stochastic cell loss and chronic immune activation (*21*–*29*). Under this view, diversity passively erodes over time independent of any intrinsic homeostatic regulatory mechanisms. However, these factors fail to explain the systematic, population-wide patterns observed in large datasets, nor do they identify specific mechanisms that mediate and constrain diversity.

Understanding what determines TCR diversity is critical because it directly impacts the immune system’s ability to recognize novel antigens. Reduced TCR diversity is linked to poor health outcomes including a greater risk of infectious disease and cancer (*20, 21, 24, 26, 30*– *32*). This association with human health highlights the urgent need to understand TCR diversity within the broader context of immune homeostasis. Identifying factors that determine TCR diversity, which may themselves be intrinsically regulated, can provide new mechanistic insight into how TCR diversity is maintained and inform interventions aimed at restoring immune competence after its decline.

Here we analyze TCR*β* repertoires from *∼*30,000 individuals and show that TCR diversity is almost entirely determined by two measurable features of the repertoire: the total number of T cells sequenced (repertoire size) and the frequency of the 1,000 most abundant clones. These two quantities independently correlate with TCR diversity and together account for 96% of its variation across individuals, including its systematic dependence on age and sex, as well as its response to Cytomegalovirus (CMV) exposure. The key finding of our analysis is that the predictive power of the frequency of abundant clones arises from the apparently coordinated nature of clonal expansion across the repertoire—a property we refer to as *intrinsic clonality*. Intrinsic clonality may be a previously unrecognized feature of the immune system subject to homeostatic control, helping to explain how TCR diversity is mediated within immune homeostasis with potentially important implications for translational research.

## Results

We analyze 30,430 TCR*β* repertoires processed under standardized protocols in a CLIA^1^certified laboratory. This cohort was sequenced as part of the T-Detect COVID test which was granted Emergency Use Authorization by the Food and Drug Administration^2^ and is the same cohort analyzed by Zahid et al. (*20*). We measure the total number of T cells sequenced, *S*, and the number of unique clonotypes (i.e., richness), *D*, and refer to these quantities as the repertoire size and TCR diversity, respectively; we interpret them as relative measures of the true underlying size and diversity of the peripheral repertoire. Both quantities follow a log-normal distribution across individuals (all logarithmic values refer to base 10). We further define *S*_1000_ as the total number of T cells derived from the 1,000 most abundant clones and *P*_1000_ as the percentage of the repertoire they comprise (i.e., *P*_1000_ = 100 *× S*_1000_*/S*).

Repertoire size is influenced by both biological factors and technical variables such as input volume and measurement uncertainty. We account for measurement error in our analysis and note that repertoire size strongly correlates with the T cell fraction (i.e., the proportion of peripheral blood mononuclear cells that are T cells; Spearman *ρ* = 0.84). Analyses using either repertoire size or T cell fraction yield consistent results, indicating that our findings are robust to the chosen metric. This supports the conclusion that the relationships we observe reflect intrinsic immune properties rather than technical artifacts.

### TCR Diversity, Repertoire Size and Clonal Expansion

TCR diversity declines with age. This decline occurs *∼*10 years earlier in males than in females, leading to pronounced differences emerging in middle age (Figure 1A). After accounting for measurement uncertainty, the peak-to-peak intrinsic biological variance in *D* for the central 90% of subjects increases by a factor of 2 to 5 (0.3 dex to 0.7 dex^3^) between the ages of 20 and 80 years, respectively (*20*). Notably, inter-individual variation in *D* exceeds the systematic effects of age, sex and CMV exposure, particularly among older individuals. Repertoire size (*S*) declines with age similarly to *D* (Figure 1B), while clonal expansion (*P*_1000_) increases from 10% to 30% between ages 20 and 80 and is consistently lower in females (Figure 1C). Furthermore, CMV-positive individuals exhibit slightly lower TCR diversity, substantially larger repertoire size and higher clonal expansion (Figures 1D–F). These findings demonstrate that repertoire size, TCR diversity and clonal expansion all depend on age, sex and CMV exposure status.

**Figure 1.**
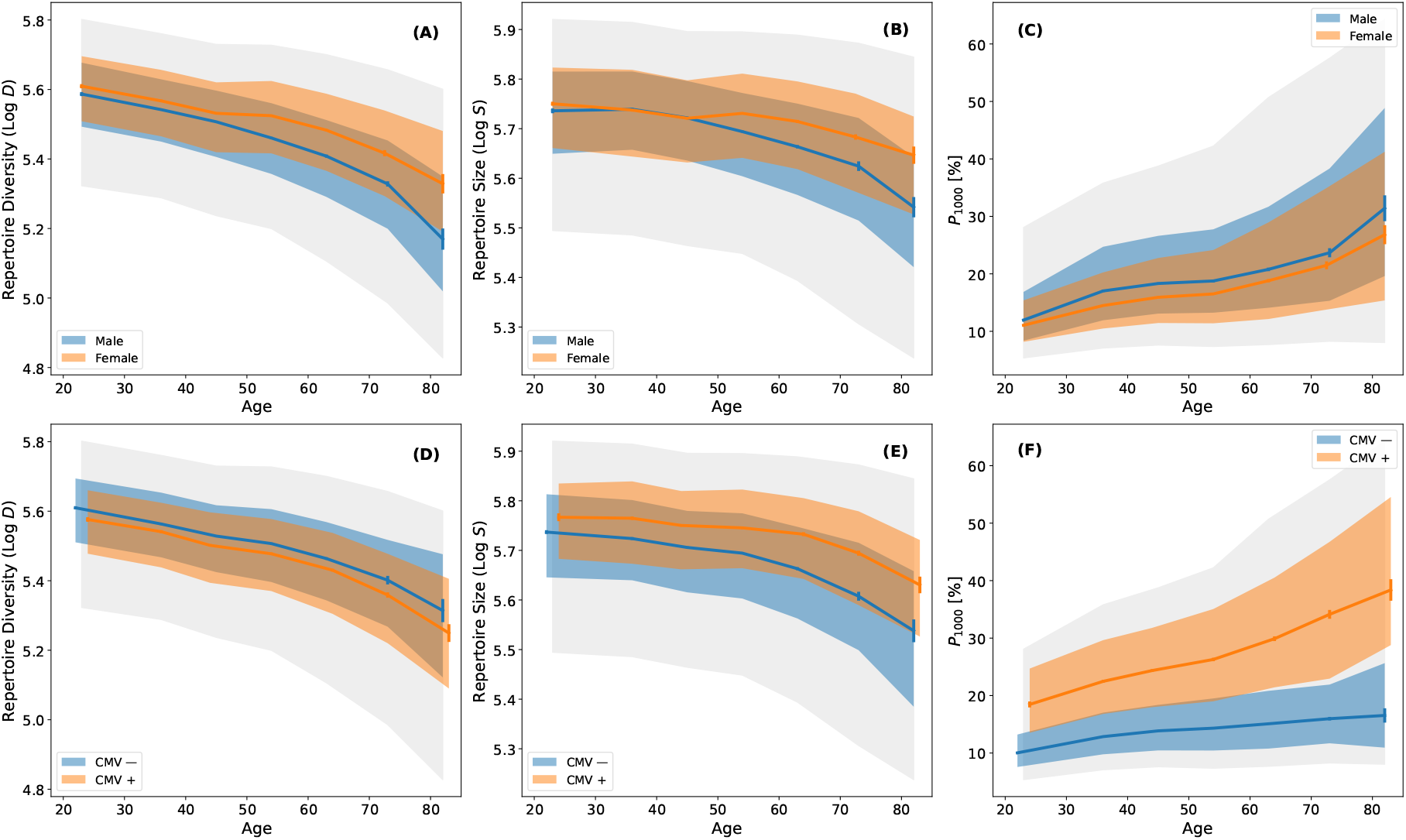
The dependence of T cell receptor diversity, repertoire size and clonal expansion on age, sex and CMV exposure status. (A) Log of TCR diversity (Log *D*) as a function of age stratified by sex. Blue and orange curves are the median diversity in decade wide age bins for males and females, respectively. Error bars are bootstrapped and blue and orange shaded regions indicate the distribution of the central 50% of the data. Gray shading represents the central 90% of the subjects. (B) Log of the total number of productive TCRs sequenced (Log *S*) in each repertoire as a function of age stratified by sex. The binning procedure and shading definitions are the same as in (A). (C) *P*_1000_ as a function of age stratified by sex. The binning procedure and shading definitions are the same as in (A). (D), (E) and (F) are the same as in (A), (B) and (C), respectively, but data are stratified by CMV exposure status rather than sex.

TCR diversity is independently correlated with both repertoire size and clonal expansion. We calculate the median *D* as a function of age, binned by *S* and *P*_1000_, respectively (Figures 2A and 2B). At all ages, individuals with high *D* tend to have low *P*_1000_ and high *S*, and vice versa. In CMV-negative individuals, larger repertoire size is accompanied by a more balanced clonal distribution, with both factors contributing to higher TCR diversity (Figure 2C). Conversely, in CMV-positive subjects, repertoire size and clonal expansion are decoupled, such that large repertoires coexist with high levels of clonal expansion. Larger repertoire sizes compensate for increased clonality in CMV-positive individuals, thereby mitigating the impact of CMV exposure on TCR diversity (*33*).

**Figure 2.**
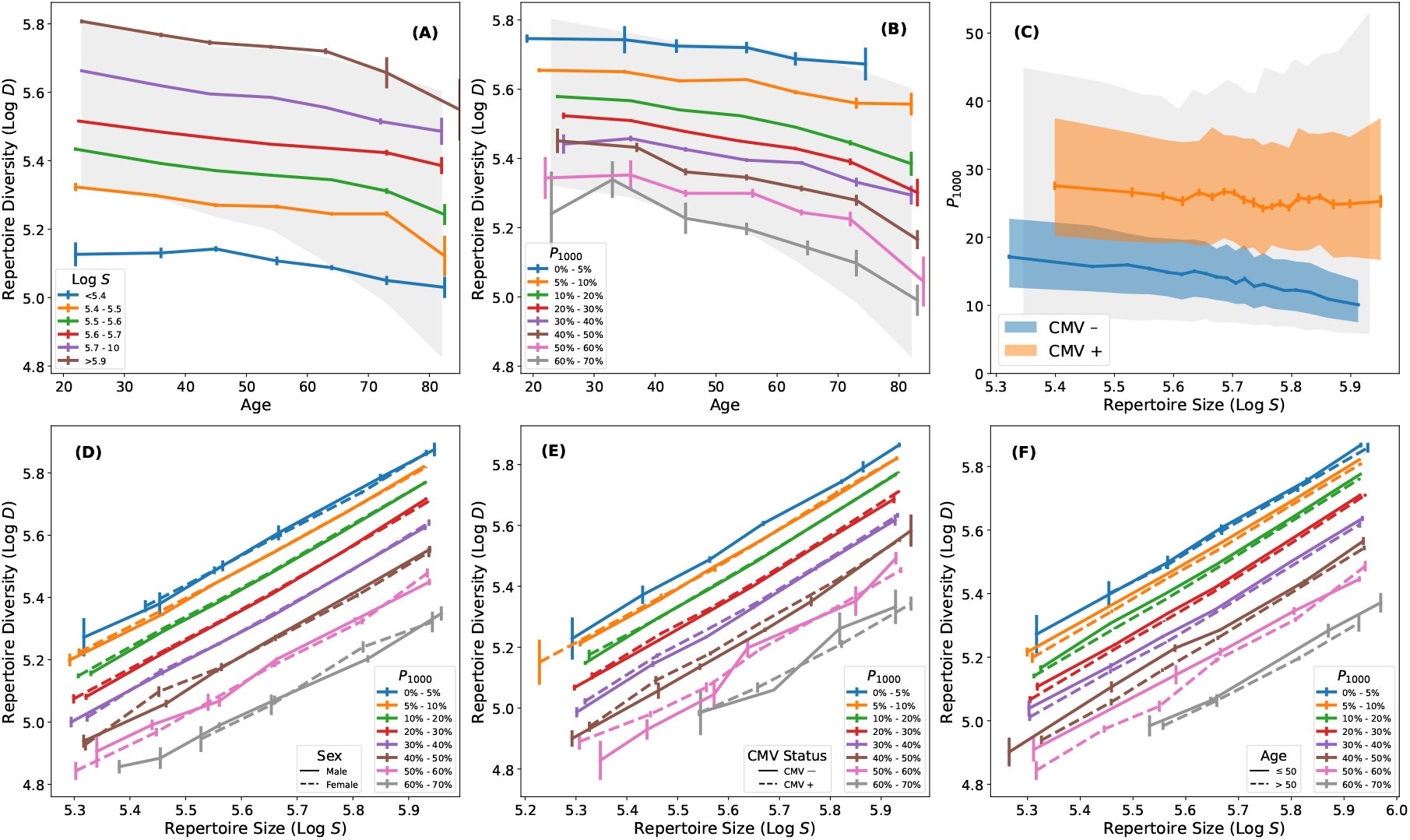
Relationship between T cell receptor diversity, repertoire size and clonal expansion. (A) Log *D* as a function of age with colored curves indicating median Log *D* in bins of Log *S*. Gray shading represents the central 90% of subjects. TCR diversity increases with repertoire size. (B) Same as in (A) but with color bars indicating median Log *D* as function of age in bins of *P*_1000_. TCR diversity decreases with increasing clonal expansion. (C) Relationship between repertoire size and *P*_1000_ stratified by CMV exposure status. For CMV-positive subjects, there is a marginal anti-correlation between *P*_1000_ and *S* (Spearman *ρ* = − 0.04, *p* = 7 × 10^*−*5^). For CMV-negative subjects, the anti-correlation is significantly stronger (Spearman *ρ* = − 0.25, *p* = 1 × 10^*−*264^). (D) Log *D* as a function of Log *S* with colored curves indicating median Log *D* in bins of *P*_1000_. The solid and dashed curves are for males and females, respectively. (E) Same as (D) but solid and dashed curves are for CMV-negative and CMV-positive subjects, respectively. (F) Same as in (D) but solid and dashed curves are subjects that are younger and older than the median age of the cohort (50 years), respectively. After accounting for repertoire size and clonal expansion, TCR diversity is independent of sex, CMV exposure status and age.

At a fixed *S, D* systematically declines with increasing *P*_1000_, demonstrating that clonal expansion reduces diversity independent of total repertoire size (Figures 2D-2F). Remarkably, after controlling for *S* and *P*_1000_, there is no residual dependence of *D* on age, sex or CMV exposure status, indicating that these biological factors influence TCR diversity through their effects on repertoire size and clonal expansion. Meaning, repertoire size and clonal expansion fully mediate the observed dependence of TCR diversity on age, sex and CMV exposure, suggesting that the relationship between these repertoire measures is independent of age, sex and CMV exposure.

Variations in repertoire size and clonal expansion almost fully account for the observed variance in TCR diversity at any given age (Figures 2A and 2B). Although *P*_1000_ increases systematically with age (Figure 1C), subjects with low *P*_1000_ and high *D* are present at all ages. The 1000 most abundant clones in any repertoire are dominated by CD8^+^ memory T cells (Figure S1), but TCR diversity (*D*) is strongly correlated with the average clonal expansion of all clones in the repertoire (Spearman *ρ* = 0.92), including naive T cells. Notably, this correlation remains significant even when the 1000 most abundant clones are excluded (Spearman *ρ* = 0.52), indicating that *P*_1000_ reflects intrinsic clonality across the repertoire.

### Modeling TCR Diversity

To better understand the factors shaping TCR diversity, we develop predictive models using biological and repertoire-derived features. We first use XGBoost, a gradient-boosted decision tree algorithm (*34*), to model TCR diversity as a function of age, sex and CMV exposure status. All three features are predictive (Figure 3A) and the model broadly captures the systematic dependence of TCR diversity on these variables (Figure 3B). Next, we include repertoire size (*S*) and clonal expansion (*S*_1000_^4^). Adding these features substantially improves model performance: *S* and *S*_1000_ together explain approximately 96% of the intrinsic variance in TCR diversity (see Materials & Methods). Furthermore, including *S* and *S*_1000_ eliminates the predictive value of sex, age and CMV status, confirming that their effects on TCR diversity are mediated through repertoire size and intrinsic clonality.

**Figure 3.**
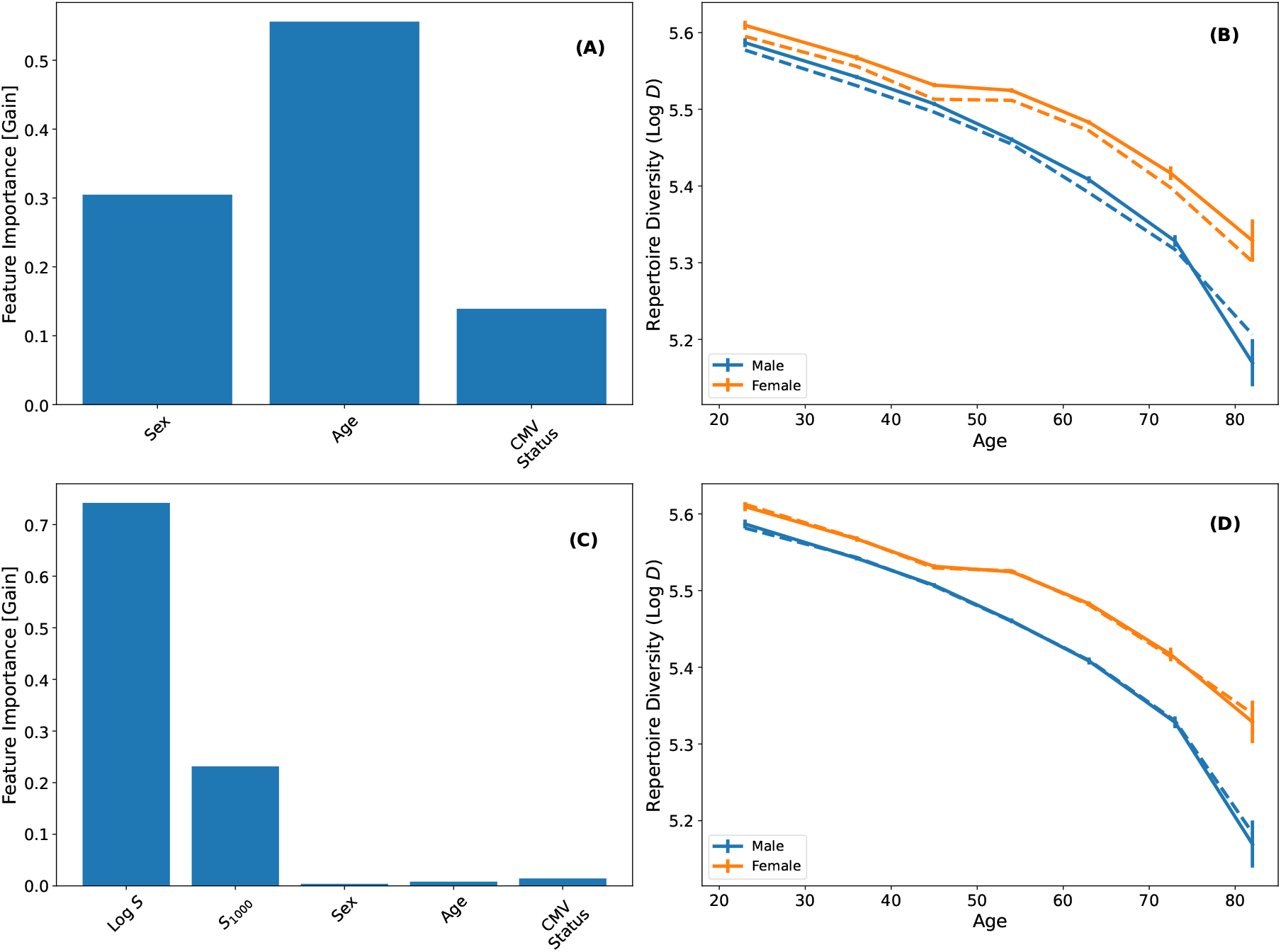
Modeling TCR diversity using XGBoost. (A) Feature importance of sex, age and CMV status in predicting TCR diversity. (B) Solid lines show the median Log *D* as a function age stratified by sex. The dashed lines show the model predictions generated via five-fold cross-validation. (C) Feature importance when including Log *S* and *S*_1000_ as features in the model. The negligible contribution of age, sex and CMV status demonstrates that repertoire size and intrinsic clonality account for the dependence of TCR diversity on these factors. (D) Same as (B) but for a model including repertoire size and intrinsic clonality as parameters. Repertoire size and intrinsic clonality robustly predict TCR diversity and are better predictors of its systematic dependence on age and sex than the model shown in (A) and (B).

The relationship between TCR diversity, repertoire size and intrinsic clonality is independent of age, sex and CMV exposure status. We fit a linear model to explicitly quantify this relationship:

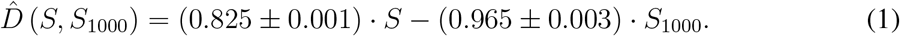

Here 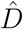 represents the predicted TCR diversity as a linear function of *S* and *S*_1000_. Equation 1 describes *D* as increasing with *S* and decreasing with *S*_1000_. Notably, the coefficient of *S*_1000_ is close to unity, indicating that TCR diversity decreases nearly one-to-one with increasing *S*_1000_. The model yields an *R*^2^ of 0.96 (Figure 4), consistent with the *∼*4% residual intrinsic scatter estimated from our XGBoost model, further supporting the robustness and completeness of the relationship. These results reinforce the idea that intrinsic clonality (captured by *S*_1000_ and *P*_1000_) reflects a systemic repertoire characteristic and is a key variable mediating TCR diversity. Despite CMV-positive and CMV-negative individuals exhibiting distinct relationships between repertoire size and intrinsic clonality (Figure 2C), both groups follow a consistent relationship linking these variables to TCR diversity, underscoring the universal and fundamental role of this relationship in characterizing immune homeostasis.

**Figure 4.**
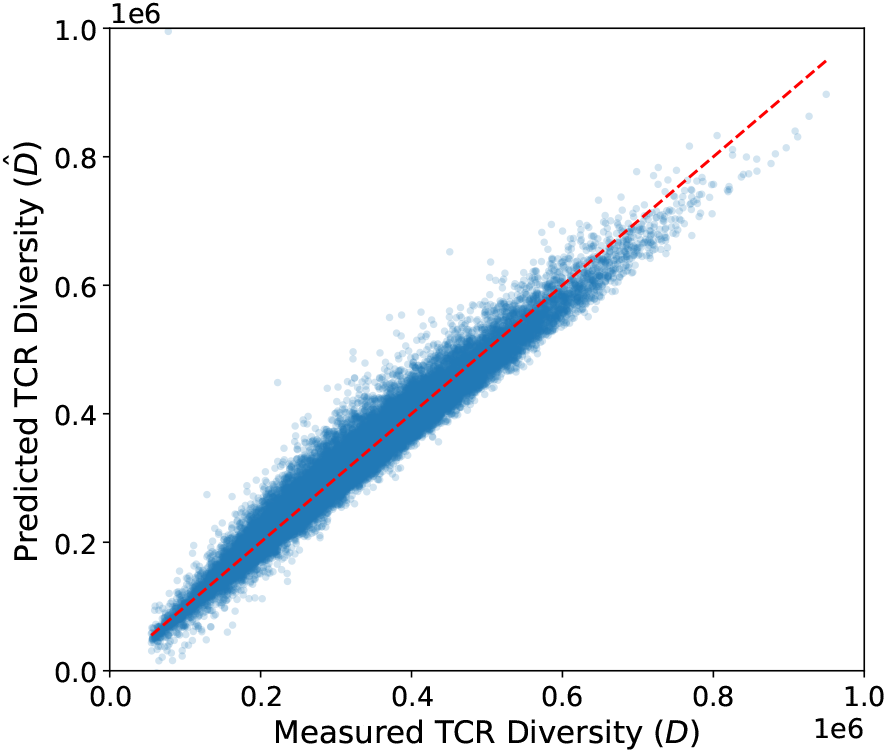
A Linear Model of TCR diversity. We model *D* as a linear function of *S* and *S*_1000_ and fit the data in five-fold cross-validation. We plot the predicted TCR diversity as a function the measured TCR diversity. The red dashed line shows one-to-one correspondence. The linear model accurately describes the measured TCR diversity (*R*^2^ = 0.96).

We note that the choice of the 1000 most abundant clones is not inherently special. For instance, when using the 10 or 100 most abundant clones (*S*_10_ or *S*_100_, respectively) as features (Figure S3), the model performance is only slightly degraded—likely due to greater statistical uncertainty in these measures. Additionally, we find that low TCR diversity does not appear to be linked to any specific or specialized set of immune exposures and HLA genotypes are not predictive (see Supplementary Material). Taken together, these results strongly suggest that repertoire size and intrinsic clonality are the primary determinants of TCR diversity.

## Discussion

TCR diversity is essential for immune competence, enabling the recognition and elimination of diverse threats. We show that repertoire size (*S*, which is strongly correlated with T cell fraction) and intrinsic clonality (*S*_1000_) fully explain the variation in TCR diversity (*D*) across individuals, including its systematic dependence on age, sex and CMV exposure. We emphasize that this relationship is not a mathematical artifact or tautology. Although *S*_1000_ is derived from a subset of highly expanded clones, it robustly predicts *D* which is a global property of the repertoire, indicating that *S*_1000_ reflects a repertoire-wide pattern of coordinated clonal expansion, i.e., intrinsic clonality. The predictive power of intrinsic clonality does not depend on selecting the 1,000 most abundant clones; using as few as 10 or 100 most abundant clones yields consistent results, reinforcing the idea that intrinsic clonality captures a fundamental biological property of the immune system. Thus, our findings reveal a simple but powerful organizing principle: TCR diversity is an emergent property of the immune system, arising from a fundamental relationship between repertoire size, intrinsic clonality and TCR diversity itself.

CMV exposure underscores the generality and resilience of the relationship between *D, S* and *S*_1000_, demonstrating that it holds even under strong immune perturbations. Unlike acute infections such as SARS-CoV-2 or other chronic herpes viruses like Epstein-Barr Virus, which have significantly smaller and more transient effects on the repertoire, CMV strongly perturbs homeostasis by increasing both repertoire size and intrinsic clonality. The chronic nature of CMV alone does not fully explain its outsized impact on repertoire structure and the biological reasons for its influence remain incompletely understood (*35*). Nevertheless, the consistency of the relationship between *D, S*, and *S*_1000_ across CMV-exposed and -unexposed individuals underscores its fundamental and robust nature.

We find no evidence that shared immune exposures or unmodeled host factors explain the relationship between *D, S*, and *S*_1000_. HLA genotype does not predict *D* and both the most abundant clones and their co-occurrence patterns vary widely across individuals, consistent with their origin from disparate immune exposures (Figure S4). Additionally, a targeted search for TCRs associated with low-diversity repertoires identified no strong candidates, further suggesting that unrecognized shared exposures are not the primary driver of diversity loss (see Supplementary Methods). However, these findings are not central to our conclusions. Rather, the key result is that *S* and *S*_1000_ together explain 96% of the variation in TCR diversity, leaving little room for additional contributors. While other variables may correlate with these quantities, repertoire size and clonality are fundamental properties of the T cell repertoire. The predictive power of *S*_1000_ reflects a coordinated pattern of clonal expansion across compartments and is robust to the specific number of clones included in the calculation (Figure S3). Notably, *S*_1000_ is dominated by memory CD8^+^ T cells (Figure S1), while *D* is shaped primarily by naive CD4^+^ T cells (*36*). The near one-to-one inverse relationship between *S*_1000_ and *D* (Equation 1) supports systemic coordination, possibly explaining the observed stability of clonal hierarchy over time (*37*). Together, these findings point to intrinsic, homeostatic regulation of the T cell repertoire rather than extrinsic factors.

The systemic coordination required to maintain the relationship between repertoire size, clonality and diversity points to regulation by intrinsic mechanisms. The naive T cell pool— the main source of TCR diversity (*25*)—is sustained by IL-7 signaling and interactions with self-peptides presented by HLA molecules (*38, 39*), suggesting a competitive feedback loop in which cytokine availability and self-recognition regulate naive T cell survival (*17, 40*). While IL-7 plays a central role in maintaining this pool, additional intrinsic signals likely contribute to broader homeostatic regulation (*41, 42*). Cytokine levels and T cell counts vary systematically with age, sex and genetic factors (*43*–*50*), suggesting genetically encoded set points that may underlie the age- and sex-associated differences in TCR diversity we observe. Moreover, genetic variation beyond HLA has been shown to influence immune traits (*44, 51*–*54*) and twin studies indicate heritability in responses to homeostatic cytokines such as IL-7 and IL-2 (*55*), as well as broader aspects of immune function (*43*). Together, these findings support the existence of intrinsic regulatory mechanisms mediated by cytokine signaling and competition for survival niches, some of which may be genetically encoded. While extrinsic factors (*56*–*59*) may modulate the system, they likely act through these intrinsic pathways.

Our analysis captures population-level trends in a cross-sectional manner. Small studies suggest TCR diversity is stable over short periods but declines with age (*60, 61*). While these studies are consistent with our finding, their small sizes underscore the need for large-scale, longitudinal studies to help establish intrinsic clonality as a repertoire-wide feature. Investigating links between our findings and immunosenescence and inflammaging (*62, 63*) may offer further insights, as the mechanisms we propose may help explain key aspects of these age-associated phenomena. Clarifying how CMV perturbs homeostasis, especially its effect on expanding repertoire size, may help identify specific regulatory mechanisms. Future studies leveraging high-throughput proteomics in large cohorts could elucidate how systemic cytokine levels influence TCR diversity and help identify molecular mediators of immune homeostasis. Continued integration of immune repertoire data with genomic profiling may help clarify how genetic variation modulates repertoire structure through the regulatory mechanisms we describe. Our findings provide a conceptual framework for investigating how intrinsic and extrinsic forces jointly regulate immune homeostasis, with implications for aging, disease susceptibility and therapeutic intervention.

T cells are essential for maintaining human health and their dysregulation contributes to a wide range of diseases, motivating therapeutic efforts to restore immune function (*64, 65*). A consistent feature of immune dysfunction is the loss of TCR diversity which is linked to poor clinical outcomes (*20, 21, 24, 26, 30*–*32*). Our findings suggest that TCR diversity is not directly regulated but instead emerges from clonal dynamics governed by repertoire size and intrinsic clonality, two properties that are likely directly regulated within immune homeostasis. By identifying these core determinants, our work provides guidance for therapeutic efforts aimed at preserving TCR diversity and restoring immune balance.

## Materials & Methods

### Sequencing of Human Samples

Details of the sequencing data and IRB information are provided in Zahid et al. (*20*), here we highlight the most salient information. The CDR3 of TCR*β* chains of T cells is sequenced with a multi-plexed PCR typically using 18*µ*g of genomic DNA (*66*–*69*). The median sequencing depth is 518,618 TCRs with 95% of subjects having a sequencing depth between 222,082 and 853,647. The median TCR diversity is 319,802 and 95% of subjects have values between 120,576 and 597,890. 95% of subjects have ages ranging between 20 and 74 years with a median age of 50 years. Sex is self-identified with males comprising 47.2% and females comprising 52.5% of subjects.

### T cell Based CMV Diagnostic

We use a sensitive and specific T cell based diagnostic on the T Detect Covid cohort to identify subjects exposed to CMV. We use a method previously described in (*70*–*72*), which statistically identifies disease associated TCRs based on serologically labeled cases and controls. 2181 labeled samples were used to build a CMV classifier with an area under the receiver operating characteristic curve (AUROC) of 0.96, measured on the same holdout set used in Emerson et al. (*70*). The performance of the T cell based test is comparable to serology and is limited by the accuracy of the serological labels. This diagnostic test allows us to identify subjects who are exposed to CMV using only their sequenced repertoire. Zahid et al. (*20*) demonstrate that CMV exposure primarily impacts TCR repertoire size.

### Fitting TCR Diversity

We first fit TCR diversity using the XGBRegressor routine implemented in version 2.1.6 of the XGBoost algorithm (*34*). We select XGBoost because of its ability to capture non-linear relationships, its strong out-of-the-box performance and its flexibility handling categorical variables like sex and CMV exposure status. We adopt the default hyperparameters of the algorithm and use its default squared error loss function. We derive predictions of TCR diversity using a five-fold cross-validation scheme implemented in the routine cross val predict from the scikit-learn (*73*) package version 1.2.0. We fit the model to a random 80% of the data and predict on the remaining 20%. This process is repeated across five distinct, randomly generated 80/20 splits of the data, ensuring that every data point is predicted without being used for model fitting. We determine feature importance by fitting all the data simultaneously.

We next fit the TCR diversity using a linear model described in Equation 1. We generate predictions in five-fold cross-validation and derive parameters by fitting all the data. We optimize the two parameters using the the optimize.curve fit module in version 1.15.2 of the SciPy package (*74*). We fit a linear model using the full dataset and generated parameter uncertainties using bootstrap resampling. Model evaluation was performed using five-fold cross-validation and Figure 4 shows predicted versus observed TCR diversity values under this validation framework. The reported coefficients and uncertainties reflect the best-fit values and 1*σ* bootstrapped error estimates.

### Estimating Residual Intrinsic Scatter

To quantify the residual intrinsic (biological) scatter in the XGBoost model’s prediction of TCR diversity, we estimate and subtract the contribution of measurement error as:

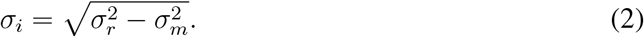

Here *σ*_*i*_ is the intrinsic biological scatter, *σ*_*r*_ is the model uncertainty and *σ*_*m*_ is the measurement uncertainty. The rationale is that for a perfect model the residual intrinsic scatter would be *σ*_*i*_ = 0, meaning the model’s uncertainty would be entirely limited by the measurement error.

Given that measurement errors in repertoire size and TCR diversity are correlated, we estimated the minimum achievable error (MAE) based on variability in *D/S*, the ratio of TCR diversity to repertoire size. Using repeat independent measurements from the same subjects, we calculated the MAE as the standard deviation of differences in Log *D/S*, yielding 0.027 dex (see Supplementary Material; Figure S2). We then fit TCR diversity using only *S* and *S*_1000_ as features and found the standard deviation of the fit residuals to be 0.031 dex. Subtracting the MAE from model uncertainty in quadrature yielded a residual scatter of 0.015 dex, indicating that approximately 4% of the intrinsic variability in TCR diversity remains unexplained by *S* and *S*_1000_.

## Data Availability

Data table with TCR repertoire metrics available at https://doi.org/10.5281/zenodo.14976210.

## Acknowledgments

We thank Ruth Taniguchi for discussion that helped improve the manuscript.

## Footnotes

1 linical Laboratory Improvement Amendments of 1988

2 https://www.fda.gov/media/146481/download

3 Here dex refers to scatter measured on a log scale such that a value of *x* dex indicates a relative difference of 10^*x*^.

4 Here we use the absolute number of T cells comprising the 1000 most abundant clones rather than their fractional representation (*P*_1000_) to avoid normalization effects that may obscure biological trends.

