## Supplementary Material for "A Fundamental Relationship between TCR Diversity, Repertoire Size and Systemic Clonal Expansion: Insights from 30,000 TCR*β* Repertoires"

### Clonal Frequencies of Sorted Repertoires

Figure S1 shows the distribution of clonal frequencies for various T cell compartments from 45 subjects, with CMV-unexposed (Figure S1A) and CMV-exposed (Figure S1B) subjects plotted separately. Repertoire sorting and sequencing details are available in Zahid et al. (*1*). For each subject we generate five repertoires: (1) $CD4^+$ naive, (2) $CD4^+$ memory, (3) $CD8^+$ naive, (2) $CD8^+$ memory and (5) an unsorted repertoire. Since sequencing depth varies across sorted repertoires due to differences in subset abundance, we plot relative clone frequencies, defined as clone count normalized to total sequencing depth. The composition of sorted T cell compartments is expected to reflect their distribution in the unsorted repertoire, which is the primary focus of our analysis. The most relevant aspects of Figure S2 for our study are that the 1000 most abundant clones are dominated by memory $CD8^+$ T cells.

### Intrinsic Variability of TCR Diversity

We estimate our measurement uncertainties as described in the main text. Figure S2 shows these uncertainties (Figure S2A) in comparison to the XGBoost model residuals (Figure S2B). The two are comparable, indicating that the model captures nearly all the intrinsic variability in TCR diversity.

### Clonal Expansion is Intrinsic

Figure S3 demonstrates that a model using $S_{100}$ and $S_{10}$ instead of $S_{1000}$ yields consistent results. As expected, model uncertainties estimated from the residuals are larger due to the higher statistical error of $S_{100}$ and $S_{10}$, which are derived from fewer clones. Nonetheless, the fact that these models remain highly predictive—even when using only the top 10 most abundant clones—supports our conclusion that clonal expansion is intrinsic. These results also indicate that the intrinsic clonality we identify can be measured with data at significantly lower sequencing depth, such as that provided by single-cell sequencing.

### The 1000 Most Abundant Clones Respond to Diverse Exposures

We hypothesize that variability in TCR diversity is not a consequence of T cells responding to a distinct or specialized set of exposures. To test this hypothesis we first attempt to identify TCRs associated with low diversity. Using CMV-negative subjects, we define cases and controls based on the top and bottom 7% of $P_{1000}$ distributions as a function of age. At each age, subjects with the highest $P_{1000}$ (lowest diversity) are cases, while those with the lowest $P_{1000}$ are controls. The 7% threshold balances statistical power with maximizing differences in $P_{1000}$. CMV-negative subjects are chosen to eliminate CMV as a confounder. Without this exclusion, our analysis identifies TCRs previously linked to CMV by May et al. (*2*). Our selection yields $\sim$1300 cases and controls. Using Fisher's Exact Test, we test all TCRs appearing in more than one repertoire and find none strongly associated with low TCR diversity, suggesting that TCR diversity loss is not driven by a specific unidentified exposure.

To demonstrate that the intrinsic clonality we measure is not a response to a specialized set of exposures, we intersect the 1000 most abundant clones in each repertoire with a large set of public TCRs, previously identified as statistically associated with specific HLAs and enriched for memory T cells (*1*). A significant fraction of these HLA-associated TCRs are specific to antigens derived from common exposures (*1*, *3*). Figure 5A shows that $\sim$10% of the 1000 most abundant clones in our repertoires are public TCRs we have previously identified as HLA-associated. We note that because a large portion of TCRs in any repertoire are rare and therefore private, the fraction of the 1000 most abundant clones responding to common antigens is likely much greater than 10%.

We further investigate whether the 1000 most abundant clones in repertoires are responding to distinct exposures using the TCR clustering results from May et al. (*3*). These authors analyze the occurrence of HLA-associated TCRs in ∼30,000 repertoires to identify hundreds of TCR clusters based on the co-occurrence of these clustered TCRs in subsets of individuals. This co-clustering pattern is expected if multiple TCRs respond to the same immune exposure but only in the subset of exposed individuals. Thus, the specific subset of TCR co-clusters present in a subject's repertoire reflects their immune history (*3*, *4*). Using serological data for eight common infectious diseases, May et al. validate distinct TCR clusters—out of the hundreds derived—as enriched for TCRs specific to these known exposures. Thus, May et al. interpret these TCR clusters as sets of TCRs, each specific to a single immune exposure. Across all repertoires, co-clustered TCRs that intersect the 1000 most abundant clones (Figure 3A) belong to a wide variety (>30) of distinct clusters (Figure 3B), suggesting that the 1000 most abundant clones in any given repertoire are responding to a diverse set of exposures which vary among individuals.

Taken together, these results support the conclusion that the large variability in TCR diversity is not driven by any special set of pathogenic exposures but instead reflects a fundamental aspect of immune homeostasis. We note that while CMV exposure significantly effects clonal expansion, a larger repertoire size compensates this expansion such that the impact of CMV exposure to TCR diversity is minimal.

### HLA is not Predictive of TCR Diversity

Using highly sensitive and specific models for HLA prediction (*1*), we infer the 145 most common class I and class II HLA alleles for each subject in our cohort. We represent each subject's genotype as a 145-element one-hot encoded vector and use this as the feature set in a random forest regression model (Scikit-learn v1.2.0). Targeting TCR diversity and evaluating performance via five-fold cross-validation, we find that HLA genotype is a poor predictor of TCR diversity. These results suggest that HLA variation does not play a primary role in determining TCR diversity.

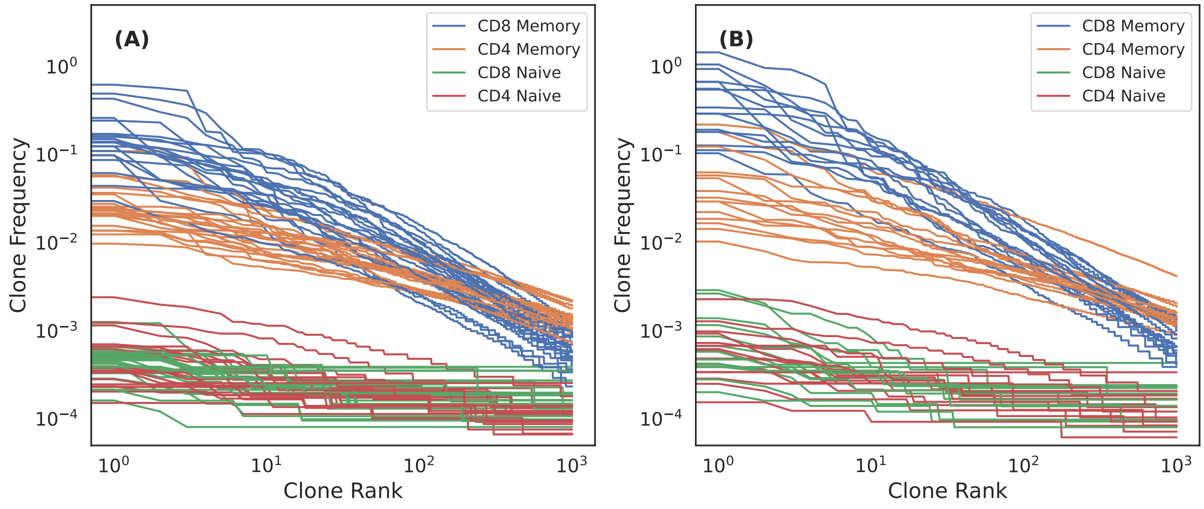


Figure S1: **Distribution of clone frequencies for various compartments.** (A) and (B) show clonal frequencies versus clone ranks derived from the sorted repertoires of 45 subjects split by CMV-negative and CMV-positive subjects, respectively. T cells are sorted on memory/naive and CD4/CD8 markers prior to sequencing. Details of the sorting procedure are provided in Zahid et al. (*1*). The 1000 most abundant clones are dominated by memory $CD8^+$ T cells.

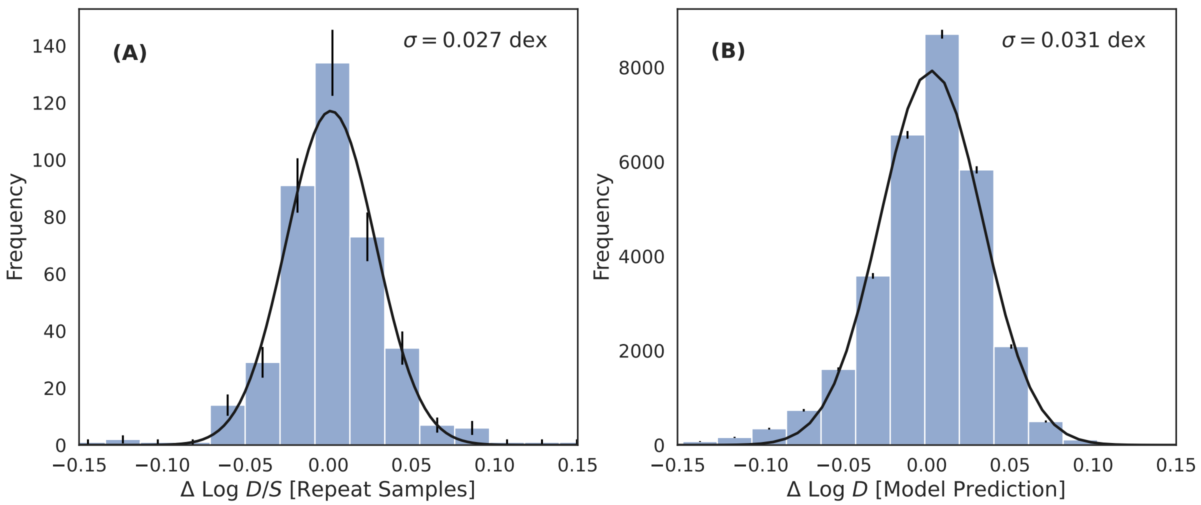


Figure S2: **Measurement uncertainty and XGBoost model residuals.** (A) Distribution of of the difference in the quantity Log $D/S$ determined from 396 repeat samples. To avoid overestimating measurement uncertainties due to a small number of outliers, we estimate the standard deviation of the distribution by fitting a Gaussian. We attribute this measure from repeat samples of the same subjects as an estimate of the measurement uncertainties and adopt the standard deviation as the minimum achievable error for any model that is not overfitting to the data. (B) Distribution of residuals of the XGBoost model fit of TCR diversity using $S$ and $S_{1000}$ as features. The model error is only slightly larger than the measurement uncertainty, indicating that the model accounts for nearly all the intrinsic biological scatter in the data.

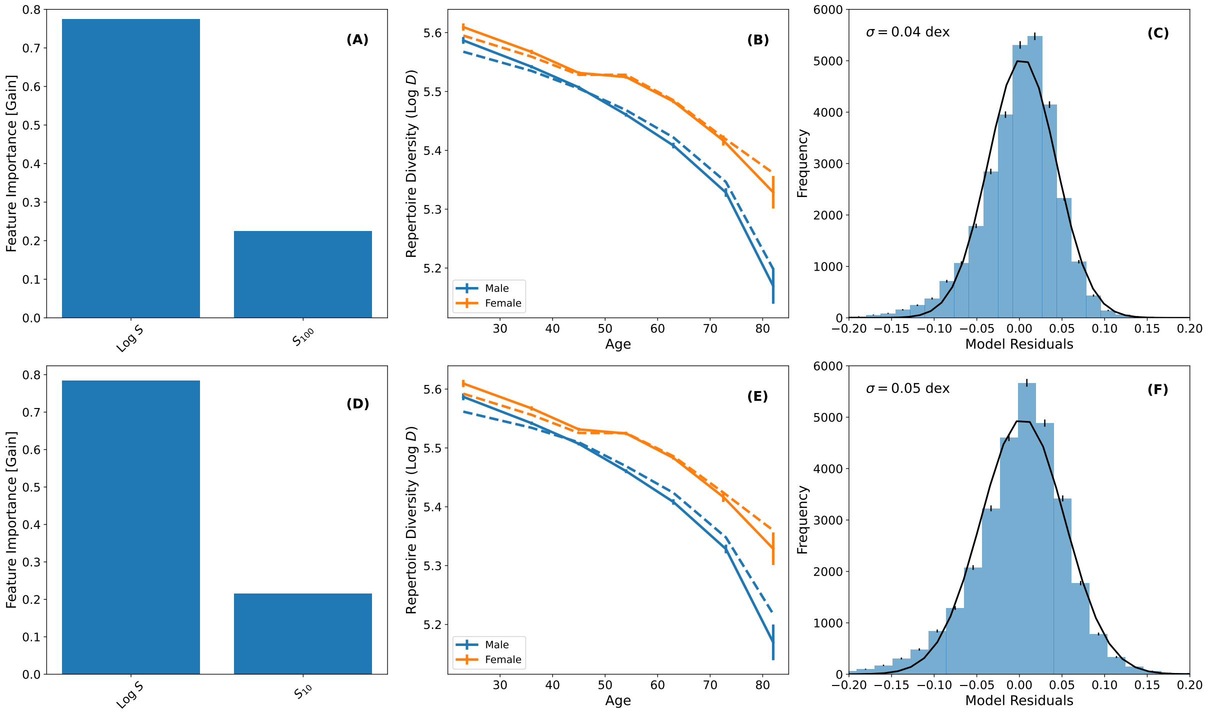


Figure S3: **Modeling TCR diversity with $S_{100}$ and $S_{10}$ instead of $S_{1000}$.** (A) and (B) are the same as Figure 2A and 2B, respectively, but for a model fit using the fraction of repertoire comprised of the 100—not 1000—most abundant clones, i.e., $S_{100}$. (C) Distribution of residuals of the model fit with $S$ and $S_{100}$. (D), (E) and (F) are the same as (A), (B) and (C), respectively, for a model using $S_{10}$ instead of $S_{100}$. Model performance is only slightly degraded when using $S_{100}$ or $S_{10}$ instead of $S_{1000}$.

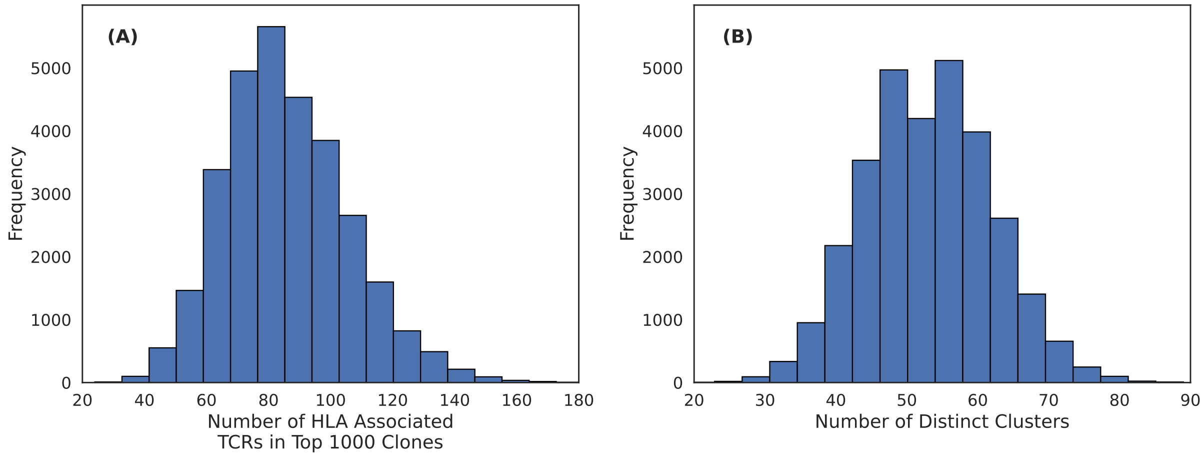


Figure S4: **Distribution of the number of TCRs in the 1000 most abundant clones that are HLA-associated and their cluster membership.** (A) Histogram of the number of TCRs that are one of the 1000 most abundant which are also HLA-associated public TCRs for all subjects. (B) Histogram of the number of distinct HLA-associated TCR clusters represented by the TCRs in (A) for all subjects. Each cluster is interpreted as mapping to a distinct immune exposure.

*4*. P. Chapfuwa *et al.*, presented at the The Thirteenth International Conference on Learning Representations, (`https://openreview.net/forum?id=wyF5vNIsO7`).
